# An intracranial dissection of human escape circuits

**DOI:** 10.1101/2024.01.16.575785

**Authors:** Haoming Zhang, Jiayu Cheng, Keyu Hu, Fengpeng Wang, Song Qi, Quanying Liu, Yi Yao, Dean Mobbs, Haiyan Wu

**Affiliations:** Centre for Cognitive and Brain Sciences and Department of Psychology, University of Macau, Macau, China; Department of Functional Neurosurgery, Xiamen Humanity Hospital, Fujian Medical University, Fuzhou, China; Section on Development and Affective Neuroscience, National Institute of Mental Health, U.S.; Department of Biomedical Engineering, Southern University of Science and Technology, Shenzhen, China; Department of Humanities and Social Sciences, California Institute of Technology, Pasadena, CA, U.S.; Computation and Neural Systems Program, California Institute of Technology, Pasadena, CA, U.S.

**Author notes:** Correspondence: Yi Yao; Haiyan Wu.

## Abstract

Predators attack at different spatiotemporal scales, spurring prey to elicit escape responses that range from simple motor reactions and strategic planning that involve more complex cognitive processes. Recent work in humans suggests that escape relies on two distinct circuits: the reactive and cognitive fear circuits. However, the specific involvement of these two circuits in different stages of human escaping remains poorly characterized. In this study, we recorded intracranial electroencephalography (iEEG) from epilepsy patients while they performed a modified flight initiation distance (FID) task. We found brain regions in the cognitive fear circuit, including the ventromedial prefrontal cortex and hippocampus, encoded the threat level during the information processing stage. The actual escaping stage, especially under rapid attack, prominently activated areas within the reactive fear circuit, including the midcingulate cortex and amygdala. Furthermore, we observed a negative correlation between the high gamma activity (HGA) of the amygdala and the HGA of the vmPFC and HPC under rapid attacks. This indicates that the amygdala may suppress the activity of the cognitive fear circuit under rapid attacks, enabling the organism to react quickly to ensure survival under the imminent threat. These findings highlight the distinct roles of the reactive and cognitive fear circuits in human escaping and provide accounts for the importance of fear in human survival decisions.

## Introduction

Naturalistic observations indicate that convergent defensive behaviors are exhibited across species [1, 2]. Across the mammalian kingdom, defensive behaviors (e.g., strategize, freeze, and flight) are closely linked to the spatiotemporal proximity between prey and predators [3, 4, 5, 6]. As part of a wider defensive system, directed escape is an important defensive behavior [7], where the prey needs to balance the risk of being killed against the loss of other survival needs including nourishment and mating opportunities associated with premature escape. To understand escaping behavior, researchers have proposed the flight initiation distance (FID) model [8, 9, 10]. This model measures the distance at which a prey chooses to escape from an approaching threat while considering the associated costs of escaping. The FID model also serves as a reliable index of threat sensitivity, enabling the study of human escape behavior and the underlying neural systems [11].

Both animal and human studies have consistently demonstrated the involvement of multiple brain regions in escaping decisions [12, 13, 14]. These brain regions can be categorized into two distinct circuits: the reactive fear circuit and the cognitive fear circuit [1, 15]. The reactive fear circuit primarily engages when the threat is close to the prey [16], encompassing the periaqueductal gray (PAG) [17, 18], hypothalamus [19], midcingulate cortex (MCC), bed nucleus of the stria terminalis (BNST) [20], and amygdala [21, 22]. Conversely, the cognitive fear circuit [23, 24], involving brain areas such as the ventromedial prefrontal cortex (vmPFC) [25, 26], hippocampus (HPC) [27, 6] and posterior cingulate cortex (PCC), is activated when the threat is not immediate. Except for the brain regions within these two circuits, the striatum and insula also play a crucial role in defensive behavior [28, 29, 30]. Despite progress in understanding the neural mechanisms of escaping in animals, and identifying the cognitive fear and reactive fear circuit in humans at various proximity threat [11, 23, 16], the underlying neural mechanisms of escaping remain unclear.

The escaping behavior encompasses various stages [31]. At its core, after the appearance of predators, prey must perceive and process threatening information (information gathering, prospection and planning), and subsequently choose the optimal moment to escape [7, 8]. However, there is still limited understanding regarding the specific roles of the two fear circuits in these distinct stages of escaping. Moreover, although the connectivity within these circuits has been investigated in the human brain [11, 23, 16], the interaction between the reactive fear and cognitive fear circuits remains unexplored, which raises another interesting question whether reactive fear circuits deactivate cognitive fear circuits under rapid attack in humans. Although there is functional MRI (fMRI) evidence of human escaping, it has not been tracked about the instantaneous neural activity throughout the entire human escaping process [32, 11]. Stereotactic electroencephalography (sEEG) allows for precise electrode placement in specific targeted deep brain structures [33, 34] and enables the simultaneous recording from both cortical and subcortical regions [35, 36, 37]. This approach enables us to acquire precise measures of local neuronal population spiking with high spatiotemporal resolution [38, 39, 40], which is critical for investigating instantaneous escaping decision-making processes. In iEEG signals, the high gamma band activity (HGA) serves as a proxy for local population activity [39], allowing us to reveal human functional network dynamics [41]. In this study, we utilized sEEG to examine the roles and interplay of HGA over the cognitive fear circuit and reactive fear circuit during two escaping phases: the threat information processing (when participants are presented with threats) and the subsequent escaping stage (when participants initiate escape).

We extend on these previous studies by proposing the following hypotheses: (i) The cognitive fear circuits are primarily engaged during the stage of processing threatening stimuli, whereas the reactive fear circuit is primarily engaged during the escaping stage. (ii) Accurate threat processing within the cognitive fear circuit is crucial for subsequent successful escapes and flexible switching of escape strategies. (iii) The activities in the reactive and cognitive fear circuits would exhibit a contradictory relationship under rapid attack, with higher activity in the reactive circuit associated with lower activity in the cognitive circuit. To test these hypotheses, we conducted sEEG recordings from 12 patients while they performed a modified FID task [11, 42] (Figure 1 A, B, C). In this task, participants were instructed to imagine themselves as prey and need to escape from the predators (Figure 1 A). There are two types of predators with different urgency: fast- vs. slow-attacking predators. We then tracked their high gamma activity (HGA) to obtain local population activity [33, 38], within the brain regions involved in the process of escaping throughout the entire escape process. We found that the neural activation in the cognitive fear circuit encoded the type of attacker. On the other hand, the reactive fear circuits primarily engaged in the process of escaping response, particularly in response to rapid attacks. Furthermore, we observed that the amygdala and the cognitive fear circuit are inversely coupled under rapid attacks. Our results provided the first human iEEG evidence for the distinct role of reactive and cognitive fear circuits in facilitating successful escaping.

**Figure 1.**
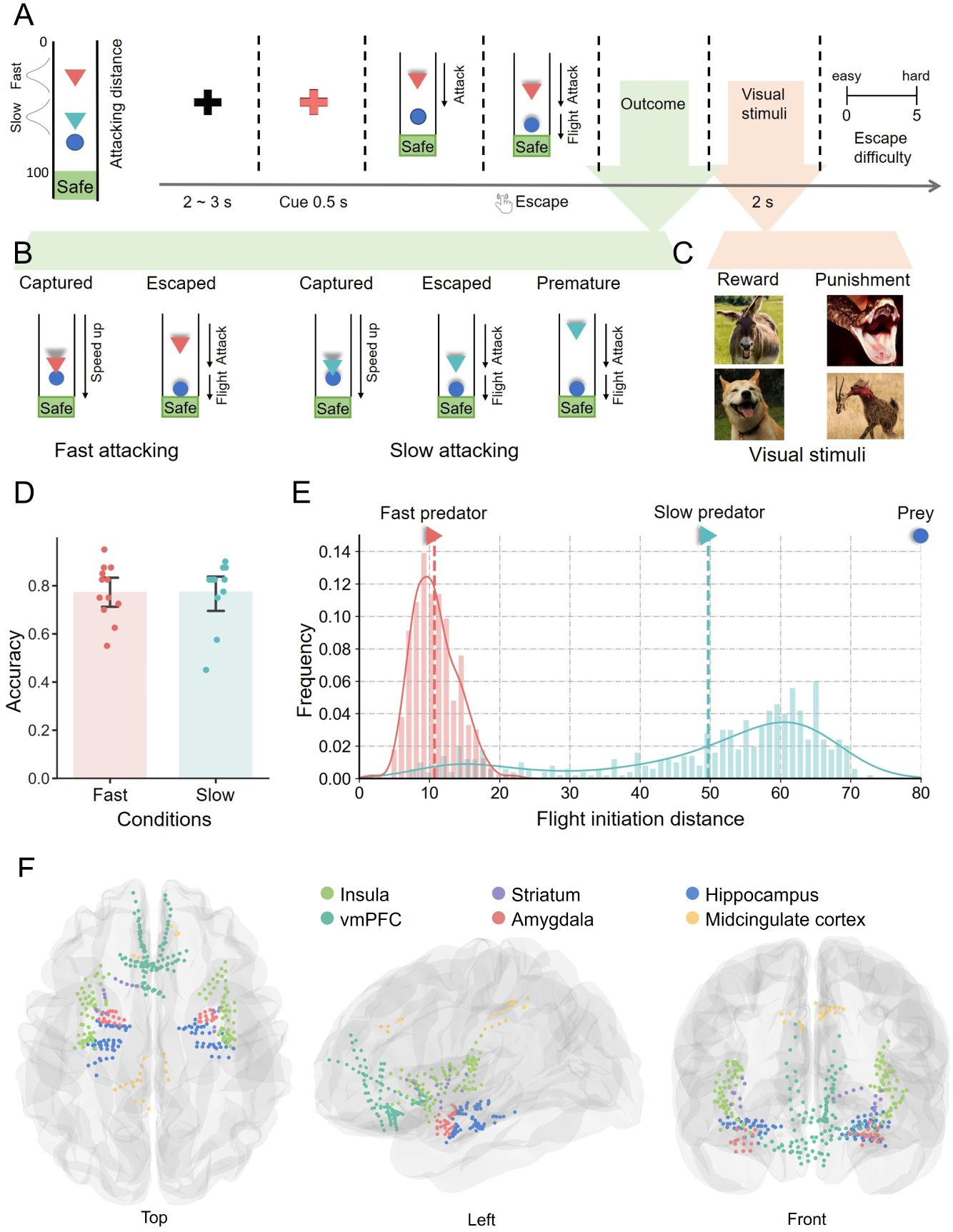
The structure of the experimental paradigm, behavioral results, and sEEG electrode recording sites. (A) The flight initiation distance paradigm. Participants (blue circle) were instructed to imagine themselves as an animal (prey) and to escape from the predator (triangle). In each trial, after the black cross baseline, participants were first displayed a colorful cue corresponding to the predator type for 0.5 seconds. Next, the predator starts to attack and participants needed to press the down arrow key on the keyboard to escape before the predator accelerated. Otherwise, the participant will be captured. To motivate longer escaping distance, participants were instructed that they needed to stay longer to get enough food under slow attack. Otherwise, premature escape will result in a lack of physical strength to escape the predator’s future capture. (B) The potential outcome of each trial. Under fast attacking conditions, the potential outcomes are captured and escaped. Under slow attacking conditions, the potential outcomes include captured, escaped, and premature escape. (C) Visual stimuli are used for reward or punishment. The reward visual stimuli used for escaped outcomes are happy images of animals. Punishing stimuli used for captured and premature conditions are terrifying images of animals or images of animals being preyed on. (D) The escaping accuracy under fast attacking and slow attacking. (E) The distribution of the flight initiation distance (FID) in response to fast attacking predator and slow attacking predator. (F) Anatomical location of all recording sites located in 6 ROIs.

## Results

### Behavioral results

We conducted intracranial recordings from 18 patients in total. Five patients were excluded from the analysis due to their limited cognitive ability or surgical history (see Method). The demographic information of the remaining 12 patients (5 females; age, 29.42 ± 10.04 years, mean ± SD) is shown in Table 1. During intracranial recordings, participants performed a modified FID task [11, 42]. In this task, participants were instructed to imagine themselves as prey and need to escape from the predator, during which they were also instructed to escape as late as possible to obtain more survival resources (Figure 1 A). The predator would initially move slowly toward the prey and then attack (accelerate) at a random distance, which varied depending on two predator types: fast-attacking predators vs. slow-attacking predators. Fast-attacking predators would initiate the attack from a greater distance, which requires the participants to make quick escape decisions. Slow-attacking predators would initiate the attack from a shorter distance, allowing for more time and a larger buffer zone to escape. In fast-attacking conditions, the potential outcomes are capture and successful escape. Under slow-attacking conditions, the potential outcomes include capture, successful escape, and premature escape (Figure 1 B). Participants will receive rewarding or punishing stimuli based on their escape success (Figure 1 C).

**Table 1.** The participant information, electrode coverage, and their related task performance. HPC: Hippocampus; PHC: Parahippocampal;

| SBJ | Age (years) | Gender | Number of Electrodes |  |  |  |  |  | Number of Correct Trials (40 in total) |  |
| --- | --- | --- | --- | --- | --- | --- | --- | --- | --- | --- |
|  |  |  | vmPFC | Striatum | HPC | Insula | Amygdala | MCC | Fast Escaped | Slow Escaped |
| S01 | 48 | F | 6 | / | 10 | 11 | 1 | 1 | 38 | 33 |
| S02 | 32 | M | 5 | / | 13 | 6 | 3 | / | 30 | 33 |
| S03 | 48 | F | / | 2 | 9 | 8 | / | / | 34 | 31 |
| S04 | 30 | M | / | / | 5 | 9 | / | / | 29 | 33 |
| S05 | 32 | F | / | / | 16 | 9 | 3 | / | 33 | 36 |
| S06 | 25 | F | 15 | 12 | 4 | 11 | 4 | / | 35 | 30 |
| S07 | 32 | M | 20 | / | 7 | 12 | 5 | / | 33 | 33 |
| S08 | 15 | M | 24 | 1 | 14 | 10 | 7 | 13 | 28 | 18 |
| S09 | 21 | F | 39 | 1 | 5 | 19 | 6 | / | 25 | 35 |
| S10 | 29 | M | / | 3 | / | 14 | 3 | 1 | 30 | 33 |
| S11 | 25 | M | / | / | / | / | / | 5 | 22 | 33 |
| S12 | 16 | M | / | / | 6 | 3 | / | 2 | 35 | 35 |
| Total |  |  | 109 | 19 | 89 | 112 | 32 | 22 |  |  |

The participants achieved an average accuracy of approximately 80% in both fast-attacking (0.775 ± 0.109; mean ± SD) and slow-attacking (0.777 ± 0.127; mean ± SD) conditions. There was no significant difference in escaping accuracy between the two conditions (*t*_12_ = 0.045, *p* = 0.965; paired t-test) (Figure 1 D). In the slow-attacking condition, the premature escape rate was 0.169 ± 0.147 (mean ± SD). Then, we examined flight initiation distance, which means when the predator chose to escape under different attacking conditions. We only focused on the not captured trials (see Figure 1 E). Under the fast-attacking condition, for all escaped trials, participants chose to escape when the predator approached a short distance (10.728 ± 3.171 units; mean ± SD). Under the slow-attacking condition, for all escaped and premature trials, participants chose to escape when the predator approached a long distance (49.708 ± 17.275 units; mean ± SD). The average predator approach distance in premature escape trials was 17.426 ± 6.669 units (mean ± SD), while the average predator approach distance in successfully escaped trials was 56.718 ± 8.838 units (mean ± SD) (see Figure S2).

### Intracerebral neural activation encodes the threat level during the information processing stage

We then examined iEEG responses in 6 brain regions within the defensive circuits: vmPFC, HPC, striatum, amygdala, MCC, and insula (Figure 1 F, Methods). To determine the role of neurons in specific ROI, we extracted the high gamma band activity (HGA) power from 70-120 Hz at each electrode in certain ROI as a proxy for local population activity [40, 43, 44] (Figure S1). We focused on the HGA during two distinct stages of escape: the threat information processing stage and the escaping stage. The threat information processing stage encompasses the period when participants begin receiving the experimental stimuli, including 0.5 seconds for attacking-type cues and 0.5 seconds for predator attacks. The escaping stage encompassed the period surrounding participants’ key presses to initiate escape, spanning 0.4 seconds before and 0.4 seconds after the escape key press. In each stage, we used a one-way ANOVA test to explore whether there were significant differences between the HGAs under different conditions. To determine whether the HGA in each condition significantly deviated from the baseline, we utilized paired t-tests. To address the issue of multiple comparisons, we conducted cluster-based permutation analysis and retained only the statistical test results within significant time clusters (*p <* 0.05).

We first examined whether there was a significant difference between the HGA under fast-attacking stimuli and slow-attacking stimuli during the threat information process stage (Figure 2). Due to the various potential causes for failed escape trials, such as errors in threat calculation or inattention, we focused our analysis exclusively on trials that achieved successful escapes. This allowed us to ensure that the included trials captured the essential neural activity associated with the required correct escape response. After the permutation test, we found that under slow-attacking stimuli, the HGA of vmPFC and HPC were significantly stronger than under fast-attacking stimuli. Region vmPFC contained three significant time clusters (Figure 2 A): cluster 1 (0.15 to 0.24 s, cue period, *p* = 0.0415*); cluster 2 (0.47 to 0.57 s, cue and predator attack periods, *p* = 0.0233*); and cluster 3 (0.71 to 1 s, predator attack period, *p* = 0.0002***). Region HPC contained two significant time clusters(Figure 2 B): cluster 1 (0 to 0.24 s, cue period, *p* = 0.0099**); and cluster 2 (0.64 to 1 s, predator attack period, *p* = 0.0078**).

**Figure 2.**
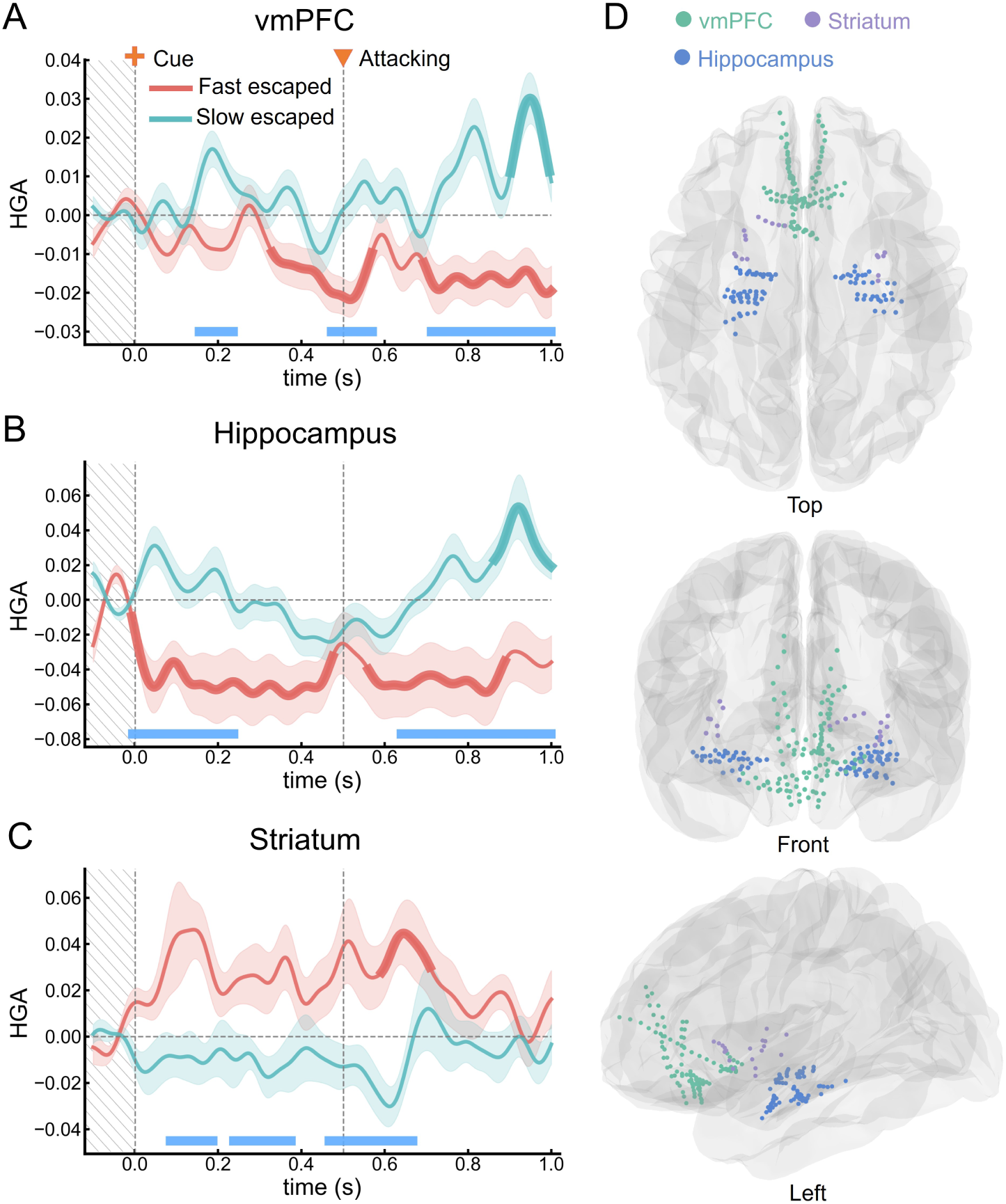
The vmPFC, HPC, and striatum encoded the threat level during information processing stage. The HGA under threat stimuli in 3 ROIs: (A) vmPFC, (B) HPC, and (C) striatum. The red and blue-green lines indicate HGA under fast escaped and slow escaped, respectively. The thicker line segment represents the time period of significant difference between HGA and the baseline. The water-blue line below indicates the time period when there is a significant difference in HGA between the two conditions. Error bars represent the standard error of the mean HGA. The stimulus (cue of attacking condition) appears at 0 s in the time series, and the attack begins at 0.5 s. (D) The anatomical location of all recording sites located in vmPFC (blue-green), HPC (blue), and striatum (purple).

On the contrary, the striatum showed significantly higher HGA under fast-attacking stimuli than under slow-attacking stimuli (see Figure 2 C). Permutation test found three significant time clusters: cluster 1 (0.08 to 0.19 s, cue period, *p* = 0.0417*); cluster 2 (0.24 to 0.38 s, cue period, *p* = 0.0407*); and cluster 3 (0.47 to 0.67 s, cue and predator attack periods, *p* = 0.0043**). The mean time-frequency spectrograms for both conditions and their contrast were plotted in Figure S3). The anatomical location of all recording sites in these 3 ROIs can be found in Figure 2 D. And the amygdala, MCC, and insula, there was no significant difference between the two conditions (Figure S4).

Besides, we also examined whether the attacking stimuli significantly influenced the HGA compared to the baseline. We found that under fast-attacking stimuli, the HGA of vmPFC (cluster 1: 0.33 to 0.57 s, *p* = 0.0021**; cluster 2: 0.69 to 1 s, *p* = 0.0012**) and HPC (cluster 1: 0 to 0.48 s, *p* = 0.0095**; cluster 2: 0.56 to 0.89 s, *p* = 0.0221*) were significantly lower than the baseline, but the HGA of striatum were significant higher than the baseline (0.59 to 0.71 s, *p* = 0.0427*). Under slow-attacking stimuli, the HGA of vmPFC (0.90 to 0.99 s, *p* = 0.0262*) and HPC (0.86 to 1 s, *p* = 0.0356*) were significantly higher than the baseline.

### Reactive fear circuit is prominently activated during the actual escaping stage

After measuring the HGA under threat stimuli, we sought to further analyze the HGA when participants chose to escape. In this analysis, we also only included trials that successfully escaped. After the permutation test, we found that the HGA during the escaping under fast-attacking was significantly higher than slow-attacking in 3 ROIs: amygdala, MCC, and insula (Figure 3). Region amygdala contained two significant time clusters (Figure 3 A): cluster 1 (−0.07 to 0.08 s, decision-making period, *p* = 0.0114*); and cluster 2 (0.14 to 0.36 s, post-decision-making period, *p* = 0.0018**). MCC contained one significant time cluster (Figure 3 B): 0.06 to 0.30 s, post-decision-making period, *p* = 0.008**). Region insula contained one significant time cluster (Figure 3 C): 0.25 to 0.38 s, post-decision-making period, *p* = 0.0311*. The mean time-frequency spectrograms for both conditions and their contrast were plotted in Figure S5). The anatomical location of all recording sites in these 3 ROIs can be found in Figure 3 D.

**Figure 3.**
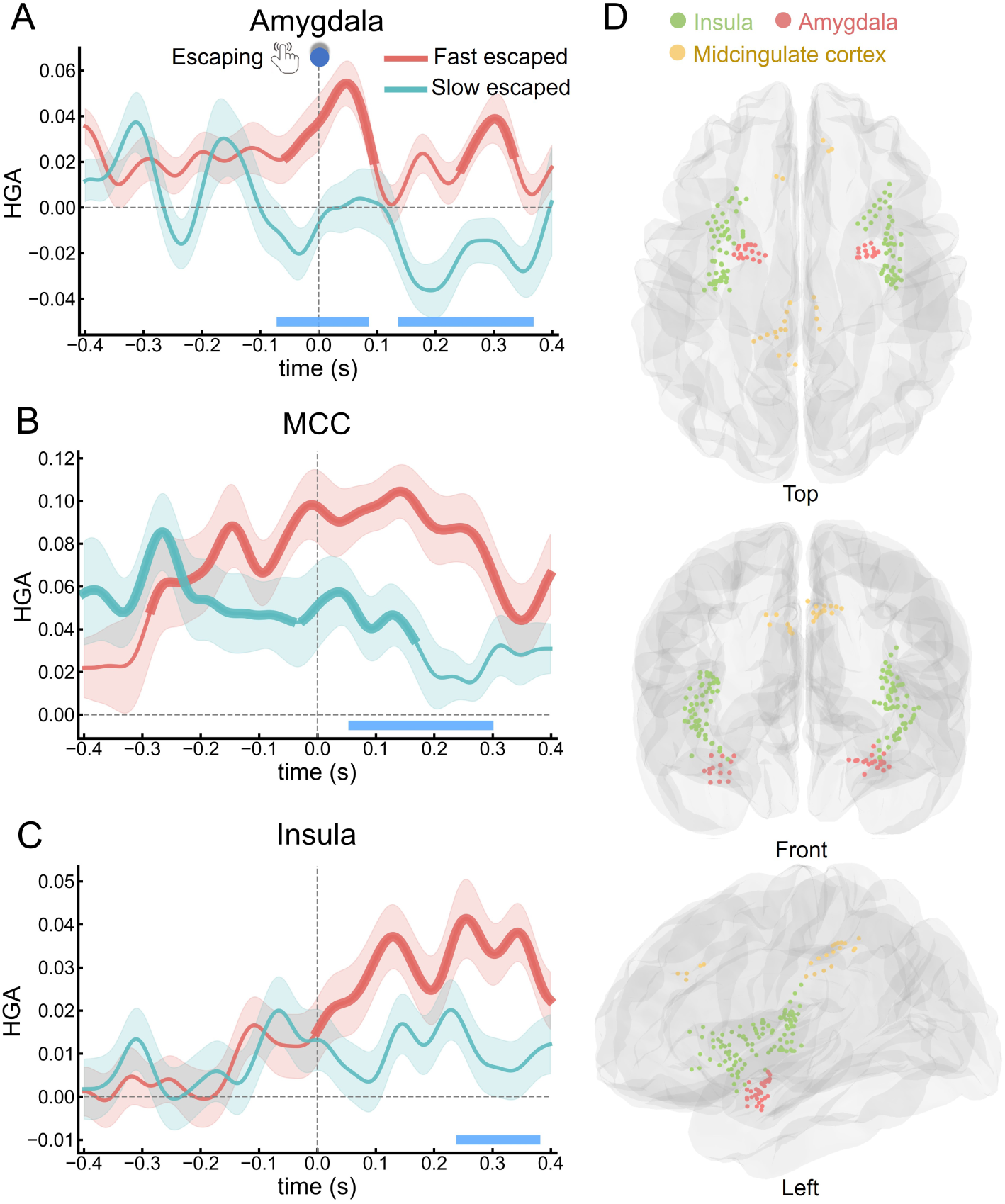
The amygdala, MCC, and insula exhibited prominent activation during escaping stage. The HGA in (A) the Amygdala, (B) MCC, and (C) the insula during the escaping stage. The red and blue-green lines indicate HGA under fast escaped and slow escaped, respectively. The thicker line segment represents the time period of significant difference between HGA and the baseline. The blue line below indicates the time period when there is a significant difference in HGA between the two conditions. Error bars represent the standard error of the mean. The zero point of the time series is the moment when the participants press the button to escape. (D) The anatomical location of all recording sites is located in insula (green), amygdala (red), and MCC (yellow).

### Cognitive fear circuit improves successful subsequent escape

After comparing the local neural activity between the conditions of fast and slow attacks, we investigated whether the neural activity involved in processing threatening stimuli would impact the subsequent success of escapes. Therefore, we compared the correct and incorrect trials under fast-attacking and slow-attacking respectively (Figure 4). We mainly focused on the specific brain regions that exhibited distinct activation patterns in response to different threat stimuli (vmPFC, HPC, and striatum). For the fast-attacking condition, we compared the HGA between the successfully escaped trials and the captured trials (Figure 4 A). We found that the HGA in the captured trials is significantly higher than the successfully escaped trials in vmPFC and HPC (Figure 4 B C), while no significant difference was found in the striatum (Figure S6). vmPFC contained three significant time clusters (Figure 4 B): cluster 1 (0.14 to 0.26 s, *p* = 0.0299*); cluster 2 (0.33 to 0.61 s, *p* = 0.0052**); and cluster 3 (0.74 to 0.85 s, *p* = 0.0115*). Region HPC contained two significant time clusters (Figure 4 C): cluster 1 (0.14 to 0.31 s, *p* = 0.0232*); and cluster 2 (0.32 to 0.48 s, *p* = 0.0237*). We also found the HGA of the vmPFC in fast-captured trials was significantly higher than the baseline (0.15 to 0.27 s, *p* = 0.0443*).

**Figure 4.**
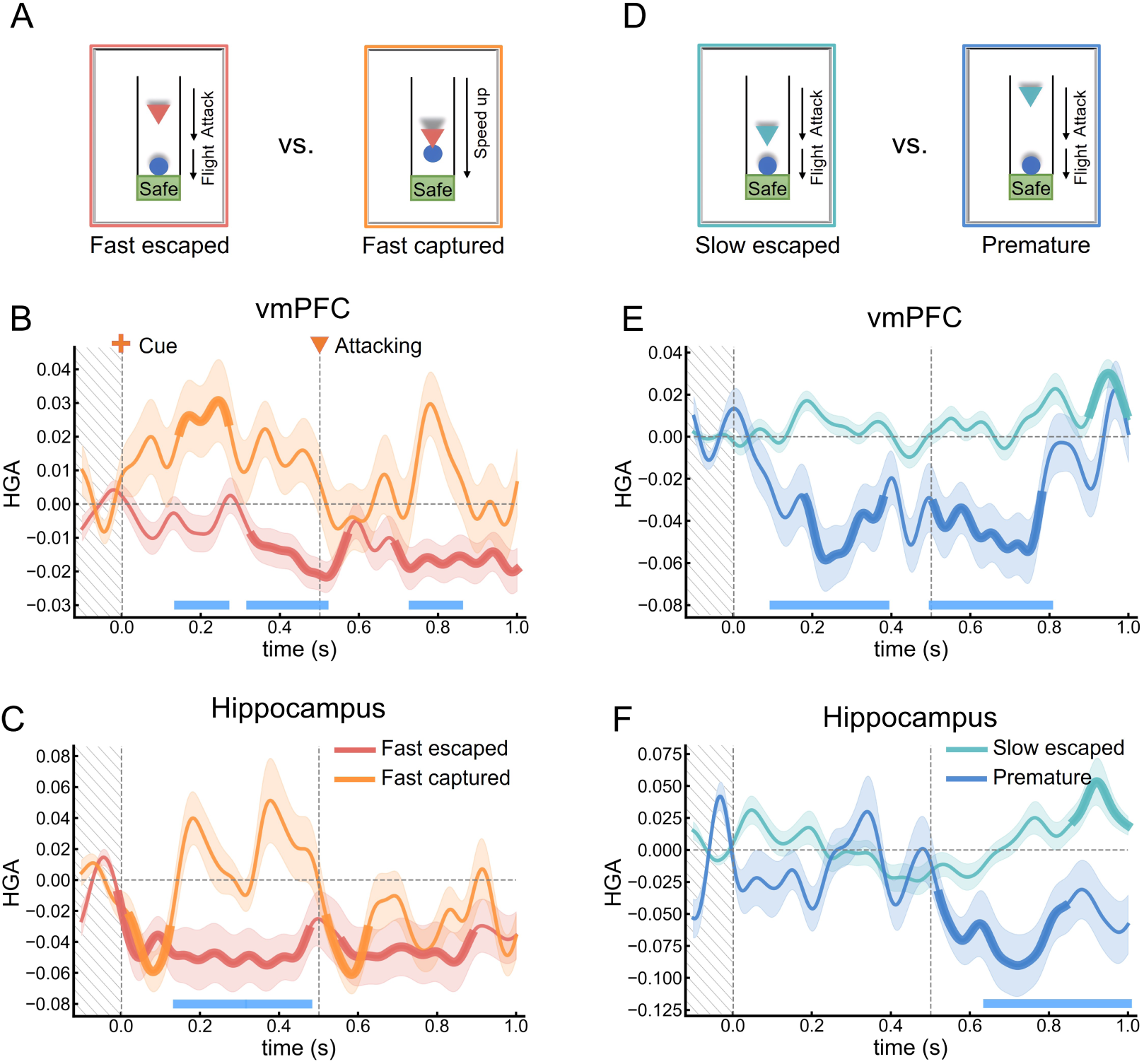
The cognitive fear circuit promotes subsequent successful escape. The HGA between (A) fast escaped trials and fast captured trials in (B) vmPFC and (C) HPC. The red and orange lines indicate HGA in the slow escaped trials and the slow premature escaped trials, respectively. The HGA between (D) slow correct escaped trials and the slow premature escaped trials in (E) vmPFC and (F) HPC. The blue-green and dark blue lines indicate HGA in the slow correct escaped trials and the slow premature escaped trials, respectively.

For the slow-attacking condition, we compared the HGA between the successful and premature escaped trials (Figure 4 D). We focused more on premature than captured trials because premature trials account for a higher proportion of failed slow-attacking trials (69.2 %). We found the HGA in the premature escaped trials was significantly higher than the successfully escaped trials in vmPFC (Figure 4 E) and HPC (Figure 4 F). In contrast, no significant difference was found in the striatum (Figure S6). Region vmPFC contained two significant time clusters (Figure 4 E): cluster 1 (0.10 to 0.39 s, *p* = 0.0004***); and cluster 2 (0.51 to 0.80 s, *p* = 0.0005***). HPC contained one significant time cluster (Figure 4 F): 0.65 to 1 s, *p* = 0.0011**. We also found the HGA of vmPFC and HPC in premature escape trials were significantly lower than the baseline (vmPFC cluster 1: 0.18 to 0.38 s, *p* = 0.0038**; vmPFC cluster 2: 0.51 to 0.78 s, *p* = 0.0012**; HPC cluster 1: 0.52 to 0.84 s, *p* = 0.0011**).

### Attacking condition switching decreases escaping accuracy

We next investigated the impact of switching between attacking conditions on both escaping behavior and its neural activation. Condition switching was defined based on the trial type (fast-attacking/slow-attacking) consistency between the previous trials (*n-1*) and current trials (*n*) (Figure 5 A). The no-switch trials consisted of two situations: fast to fast and slow to slow. On the other hand, the switch trials comprised two situations: slow to fast and fast to slow, where there was a change in the attacking condition.

**Figure 5.**
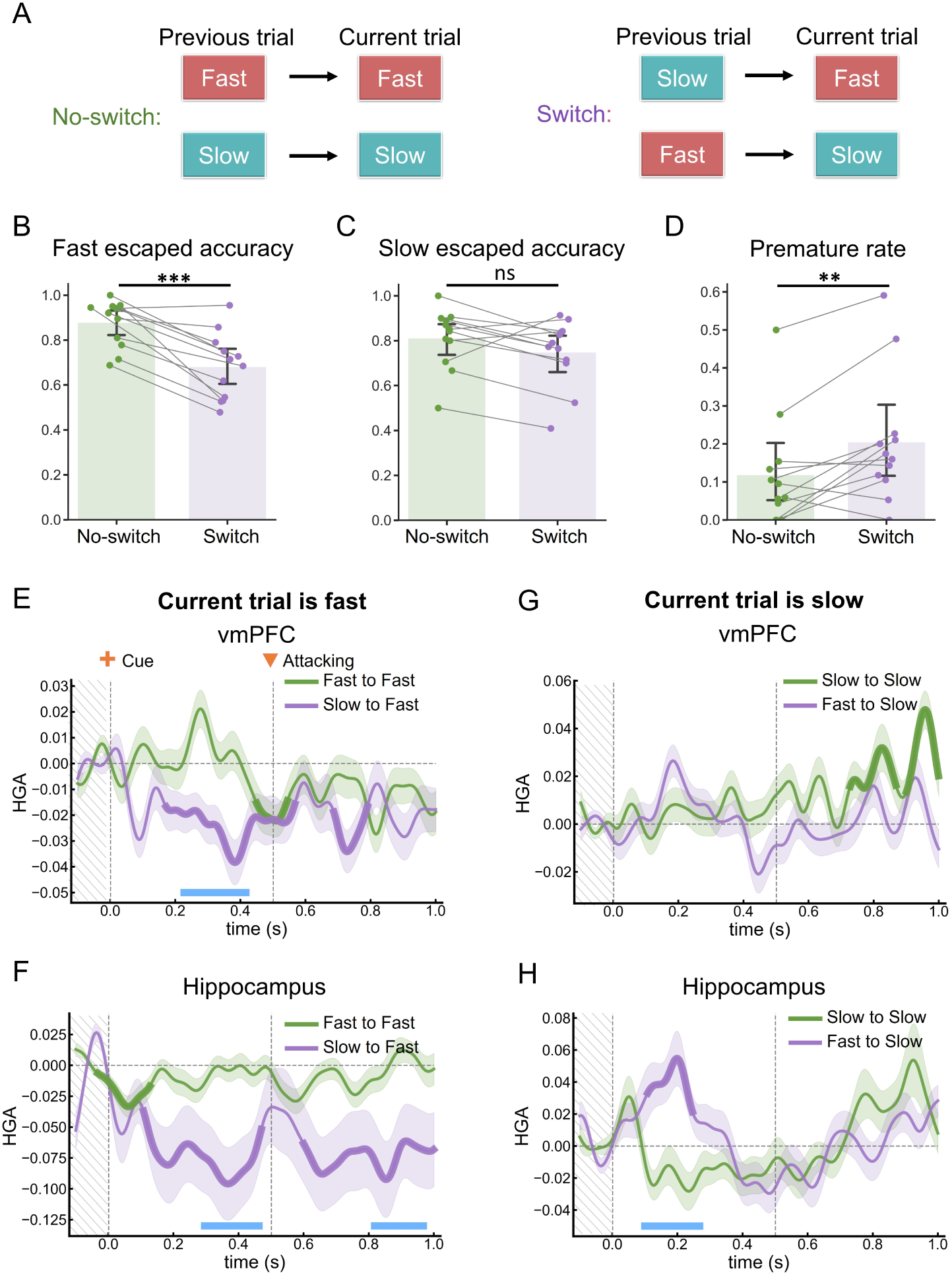
Attacking condition switching decreased escaping accuracy, while the cognitive fear circuit facilitates flexible switching. (A) Definition of escaping condition switching. (B) Escaping accuracy under fast attack in No-switch trials and switch trials. (C) Escaping accuracy under slow attack in No-switch trials and switch trials. (D) Premature rate under slow attack in No-switch trials and switch trials. Significance levels (t-test) are indicated by asterisks: \*\*\**p ≤* 0.001; \*\**p ≤* 0.01; and ns = non-significant. The HGA between the fast-fast trials (green line) and slow-fast trials (purple line) in the (E) vmPFC and (F) HPC. The HGA between slow-slow (green line) trials and fast-slow (purple line) trials in the (G) vmPFC and (H) HPC. The thicker line segment represents the time window of significant difference between HGA and the baseline. The water-blue line below indicates the time window when there is a significant difference in HGA between the two conditions. Error bars represent the standard error of the mean HGA. The stimulus (cue of attacking condition) appears at 0 s in the time series, and the attack begins at 0.5 s.

To assess whether attacking condition switching modulated escaping performance, we contrasted escaping accuracy between switch- and no-switch-trials. Under the fast-attacking, participants’ accuracy in the switching trials (0.681 ± 0.140; mean ± SD) was significantly lower (Figure 5 B; *t*_12_ = *−*5.419, *p* = 0.0002***; paired t-test) than the no-switching trials (0.878 ± 0.099; mean ± SD). Under the slow-attacking, there was no significant difference (Figure 5 C; *t*_12_ = *−*1.990, *p* = 0.072; paired t-test) between the accuracy in the switching trials (0.749 ± 0.142; mean ± SD) and the no-switch trials (0.811 ± 0.127; mean ± SD). We further examined whether the premature escaping rate will show a significant difference between the two conditions and found the premature escaping rate in the switching trials (0.119 ± 0.138; mean ± SD) was significantly higher (Figure 5 D; *t*_12_ = 3.224, *p* = 0.008**; paired t-test) than the no-switching trials (0.205 ± 0.161; mean ± SD).

### Cognitive fear circuit facilitates flexible switching between different escape strategies

After evaluating escaping accuracy, we investigated whether there were differences in the high gamma activity (HGA) between the switching condition and the no-switch condition. We focused on the vmPFC, HPC, and striatum. In the fast-attacking trials that switched from the slow-attacking trials (slow to fast), we observed a significant decrease in HGA in the vmPFC and HPC compared to the no-switch trials (fast to fast). The vmPFC region exhibited one significant time cluster (Figure 5 E): 0.23 to 0.42 s, *p* = 0.0027**. The HPC region showed two significant time clusters (Figure 5 F): 0.29 to 0.47 s, *p* = 0.0322; 0.82 to 0.97 s, *p* = 0.0469. In the slow-attacking trials that switched from the fast-attacking trials (fast to slow), we found significantly higher HGA in the HPC compared to the no-switch trials (slow to slow) (Figure 5 H): 0.10 to 0.27 s, *p* = 0.0088**. However, no significant difference in HGA was observed in the vmPFC (Figure 5 G). For the striatum, there was no significant difference between the switching trials and no-switching trials (Figure S7)

### The functional connectivity between vmPFC and HPC during attacking information processing

Next, we aimed to explore the directed FC among several crucial regions involved in human escaping. We computed the directed FC between all pairs of electrodes across different brain lobes. To obtain the directed FC, we used the time-lagged cross-correlation analysis, as described in [45], with a lag range of −300 to 300 ms. Subsequently, we examined the directed FC within the contexts of fast-attacking and slow-attacking scenarios, both at the average level across all electrode pairs and at the individual electrode pair level. At the average level, we used the significance test and cluster-based permutation analysis to explore the significant differences between the FCs in different conditions or whether the FC was different from the baseline. At the individual level, we examined whether the FC of each electrode pair significantly differed from the baseline, and recorded the time lag and the FC value at the peak.

We initially examined the directed FC between the vmPFC and HPC under attacking stimuli (Figure 6 A, B). We extracted directed FC from a total of 856 electrode pairs in 6 patients who underwent simultaneous implantation of electrodes in the vmPFC and HPC (The number of electrode pairs for each patient can be found in Table S1). At the average level (Figure 6 A, The FCs in the fast-attacking condition (−0.16 to 0.24 s, *p* = 0.0001***) and the slow-attacking condition (−0.12 to 0.12 s, *p* = 0.0001***) were both significantly higher than baseline with a peak lag of 0 s. This indicated that there was no apparent directionality in the FC between the vmPFC and HPC in both conditions. Furthermore, The FC in the slow-attacking condition was significantly higher than that in the fast-attacking condition (−0.07 to 0.09 s, *p* = 0.0001***). At the individual electrode pairs level, the FC of 65 and 95 electrode pairs showed significant increases compared to the baseline under fast attack and slow attack conditions, respectively. Conversely, the FC of 14 and 12 electrode pairs exhibited significant decreases compared to the baseline under fast attack and slow attack conditions, respectively. The peak value, corresponding peak time lags, and anatomical location can be found in Figure 6 B.

**Figure 6.**
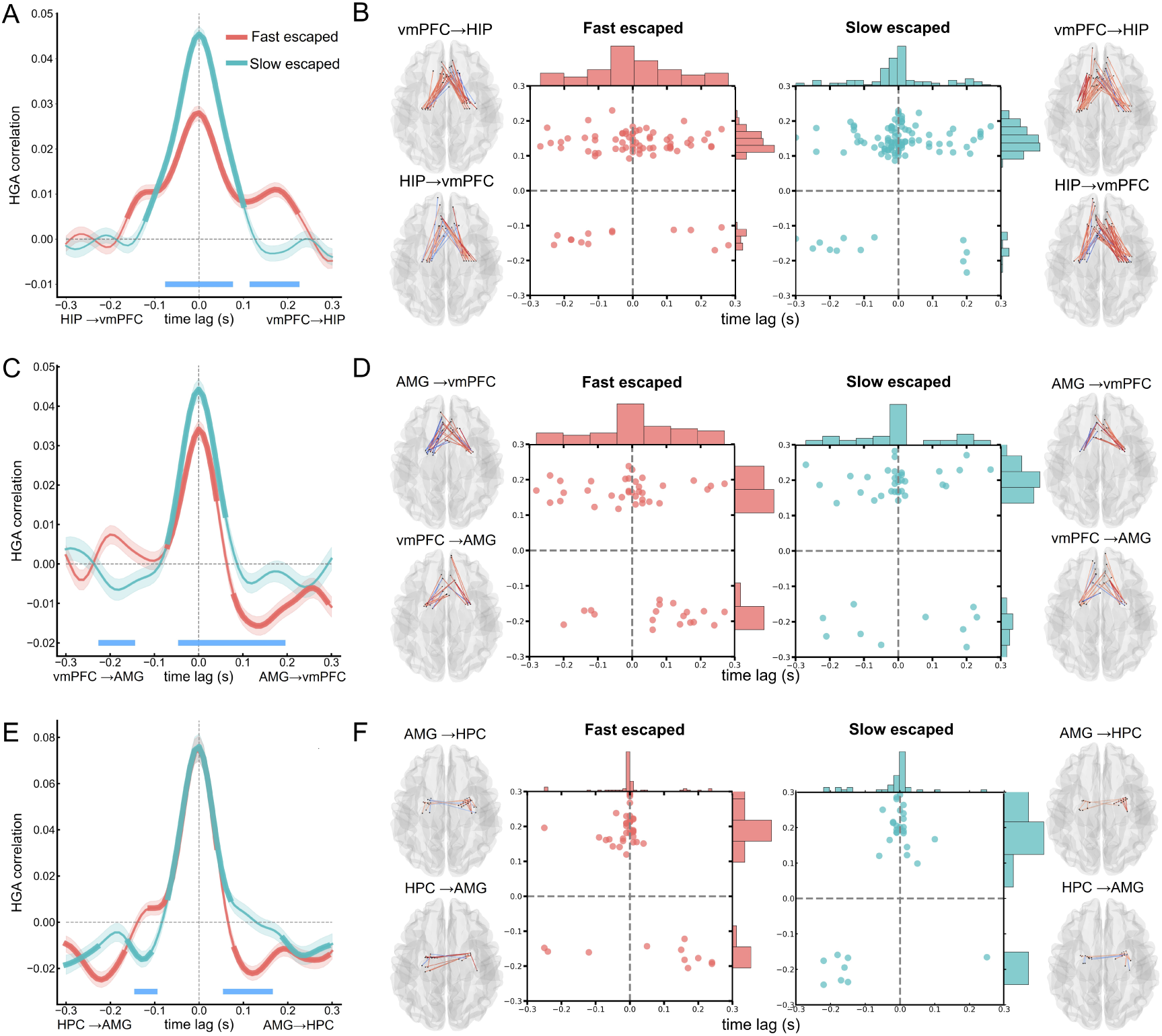
Directed functional connectivity in human escaping. (A) Average-level directed functional connectivity (FC) between the vmPFC and HPC under attacking stimuli. Positive time lags indicate vmPFC activity preceding HPC activity, while negative lags indicate the opposite. The thicker line segment represents the time period of significant difference in FC compared to the baseline. The water-blue line below indicates the time period with significant differences in FC between the two conditions. Error bars represent the standard error of the mean FC. (B) Individual-level directed FC between the vmPFC and HPC under attacking stimuli. Each point represents an electrode pair with significant FC compared to the baseline. The x-axis and y-axis of each point denote the peak FC value and the corresponding peak time lags of the significant electrode pair. The anatomical location of significant channel pairs is indicated for both directions of connectivity. (C) Average-level directed FC between the amygdala and vmPFC during the escaping decision stage. (D) Individual-level directed FC between the amygdala and vmPFC during the escaping decision stage. (E) Average-level directed FC between the amygdala and HPC during the escaping decision stage. (F) Individual-level directed FC between the amygdala and HPC during the escaping decision stage.

### The amygdala and cognitive fear circuit are inversely coupled under fast attacks

Finally, we explored whether the reactive fear circuit and the cognitive fear circuit are inversely coupled during the fast attack. We evaluated the functional connectivity (FC) between the amygdala and brain regions within the cognitive-fear circuits (vmPFC and HPC) during the escaping decision stage. We extracted directed FCs between the amygdala and vmPFC from a total of 583 electrode pairs in 6 patients (Table S1). At the average level (Figure 6 C, the FC in the fast-attacking condition (−0.07 to 0.06 s, *p* = 0.0001***) and the slow-attacking condition (−0.07 to 0.07 s, *p* = 0.0001***) were both significantly higher than baseline with a peak lag of 0 s. Besides, FC between the amygdala and vmPFC in the fast-attacking condition was also significantly lower than the baseline with a peak lag of 0.13 s (0.08 to 0.30 s, *p* = 0.0001***), indicating the activation in the amygdala deactivated the activation in vmPFC. Furthermore, the FC in the slow-attacking condition was significantly higher than that in the fast-attacking condition (−0.04 to 0.20 s, *p* = 0.0001***). At the individual electrode pairs level, we observed significant increases in FC compared to the baseline in 35 electrode pairs during fast attacks and 35 electrode pairs during slow attacks. Conversely, 18 electrode pairs showed significant decreases in FC compared to the baseline during fast attacks. Among these electrode pairs, 10 had a peak lag greater than 0 s (from amygdala to vmPFC) (Figure 6 D).

We next extracted directed FCs between the amygdala and HPC from a total of 273 electrode pairs in 7 patients (Table S1). Except that the FCs in both conditions were significantly higher than baseline with a peak lag of 0 s. We also found the FC in the fast-attacking condition was significantly lower than the baseline (0.07 to 0.30 s, *p* = 0.0001***), and also significantly lower than the slow-attacking condition (0.06 to 0.18 s, *p* = 0.0001***) (Figure 6 E). At the individual electrode pairs level, we observed significant increases in FC compared to the baseline in 45 electrode pairs during fast attacks and 33 electrode pairs during slow attacks. 11 electrode pairs showed significant decreases in FC compared to the baseline during fast attacks. Among these electrode pairs, 8 had a peak lag greater than 0 s (from amygdala to HPC) (Figure 6 F).

## Discussion

Our study utilized sEEG for the first time to investigate the defensive survival circuit in the human escaping brain. We aimed to understand how the cognitive fear circuit and reactive fear circuit are activated during specific stages of human escaping, such as the threat information processing stage and the escaping stage. In line with previous research [11, 5], we found that reactive fear circuitry shows greater activation during rapid attacks, while the cognitive fear circuit is more active during gradual attacks. Our findings demonstrated that the cognitive fear circuit primarily engages in the threat information processing stage, while the reactive fear circuit is mainly involved in the escaping stage. Furthermore, we observed a negative correlation between the HGA of the amygdala and the HGA of the vmPFC and HPC under rapid attacks. These results emphasized the dissociated and collaborative role of both fear circuits in facilitating human escape from different types of threats.

### Reactive and cognitive fear circuits exhibit greater activation under rapid and gradual attacks, respectively

Our analysis demonstrated increased neural activity in the brain regions involved in reactive-fear circuits, such as the amygdala and the MCC, under rapid attack compared to gradual attack. This finding is consistent with previous human escaping studies [11, 16]. These brain regions primarily support model-free computation and are activated when the threat is nearby [5]. Conversely, our result also revealed higher neural activation in brain regions implicated in cognitive-fear circuits, such as the vmPFC and the HPC, during gradual attacks compared to rapid attacks. These brain regions primarily support model-based planning and are activated when the threat is far away [5]. Previous research has also suggested that larger buffer distances with the predator are associated with increased activity in the cognitive fear circuit [12]. In contrast, as the buffering distance decreases, there is a shift towards increased activity in the reactive fear circuit [46, 47]. Additionally, the striatum and insula exhibit stronger neural activity under rapid attacks than gradual attacks in our results. This could be explained by the previous human fMRI studies that these two regions are associated with risk signals [48, 49, 50] and may play an important role in threat evaluation [5].

### Cognitive fear circuitry mainly functions in threat information processing stage

During the threat information processing stage, we reveal distinct neural activation patterns in the vmPFC, HPC, and striatum between rapid attacks and gradual attacks. vmPFC and HPC are significantly activated under gradual attack during the attacking period (0.5 - 1 s), further stressing the importance of cognitive circuits in threat detection. As a critical part of cognitive control, the vmPFC may play a role in optimizing goal-directed escaping behavior versus spontaneous immediate escape behavior under gradual attacks [51, 52, 53]. HPC may play a role in utilizing past threat encounters memory to process current threat information and optimize escaping behavior [54, 55]. This interpretation complements previous findings demonstrating that HPC is associated with regulating avoidance behaviors in anxiety-inducing environments [56]. We also observe that the activity of vmPFC and HPC is significantly inhibited during both the cue period (0 - 0.5 s) and the attacking period under rapid attack. Previous studies using multivoxel pattern analysis have demonstrated a stable decrease in vmPFC activity in response to fear [57, 58] and threatening stimuli [59]. Our findings expand on these results by demonstrating that urgent threats under rapid attack may also inhibit the activation of the frontal lobe and the HPC. One possible explanation is that inhibiting these model-based brain regions could recruit the shortcut of the reactive system, facilitating more effective responses to emergencies.

### Reactive fear circuitry is primarily involved during the escaping stage

During the escaping stage, our results indicate that the amygdala, MCC, and insula exhibit higher neural activation under rapid attacks compared to gradual attacks. Our findings indicate that the amygdala is involved during both the decision-making period and the post-decision-making period (outcome) under rapid attacks. It is known to be a key component involved in emotional processing and stress response when facing negative stimuli [60, 61], and plays a key role in modulating aversive behavior [62, 63]. Our analysis reveals its role in encoding escape decision-making and escape outcomes. Additionally, our findings demonstrate the MCC shows significant activation under both rapid and slow attacks. Meta-analysis indicates that the MCC consistently shows activation in response to negative affect, pain, and cognitive control [64, 65]. MCC is also a critical brain region in humans escaping under rapid attack [11]. Our study extends previous understanding and suggests that the MCC could serve as a promising brain region for carrying out aversively motivated instrumental motor behaviors under different levels of urgency [5]. Lastly, the insula is significantly activated after making escape decisions under rapid attacks, which may be associated with threat anticipation or prediction. It plays a role in encoding anticipated ”risk predictions” during risky decision-making and reflects subsequent ”risk prediction errors” when outcomes are revealed [48, 66, 67]. In our setting, there are either slow or fast threats, which means the insula may stay activated to predict the threat level. Our findings may indicate stronger threat prediction in the insula after a rapid attack, and provide evidence that such risk prediction by the insula mainly exists under more urgent threats.

### Cognitive fear circuitry supports effective escaping and strategy switching

Our findings demonstrate the crucial role of the vmPFC and HPC in guiding accurate escape decisions. While in successfully escaped trials, the cognitive fear circuit is activated under gradual attack and inhibited under rapid attack. However, in captured trials under rapid attack, the cognitive fear circuit failed to inhibit. Further, in premature trials under gradual attack, the circuit failed to activate. These results together emphasize the importance of the cognitive fear circuit in processing threat information and promoting successful human escape in both rapid and gradual attack scenarios [52, 68, 69]. Critically, the present findings suggest the key role of the vmPFC and HPC in flexible escape strategy switching under different attack conditions. The prefrontal cortex (PFC) plays a crucial role in flexible strategy selection [25] and adaptive goal-directed behavior. [70, 71, 35, 72]. Previous studies have also demonstrated the role of the HPC and PFC in adapting to novel situations and overcoming an established strategy [73]. Our analysis reveals an inconsistent neural activation between switching from rapid to gradual attacks and from gradual to rapid attacks. Compared to the no-switching fast trials, the vmPFC and HPC both show increased inhibition in the switched rapid trials. Conversely, the HPC exhibits more activation in the switched gradual trials. The cognitive fear system may guide a more flexible strategy, with sustained inhibition in fast trials, and more engagement in slow trials. An alternative interpretation of this phenomenon is that predator condition switching does not lead to consistent enhancement of neural activity. Instead, it may further polarize the original neural activity, resulting in increased inhibition under rapid attack and increased activation under gradual attack.

### Connectivity between the vmPFC, HPC, and amygdala in escaping

The synchrony between the PFC and the HPC has been consistently associated with various cognitive functions, including memory, learning, and defensive behavior [68, 73, 74, 75]. Previous rodent studies have demonstrated the role of this neural circuitry in anxiety-related responses [76, 77]. In human studies, it has been found that gradual threats increased the coupling between HPC and mPFC [16]. Moreover, the strength of FC between the HPC and the mPFC in escaping from the gradual attack correlated with the degree of trait anxiety [23]. These findings highlight the significance of the PFC-HPC connectivity in anxiety-related responses. By analyzing the FC between the vmPFC and HPC, our analyses reveals a higher level of co-activation between these regions under gradual attacks. Notably, this co-activation did not show significant directionality, suggesting a balanced and bidirectional interaction between the vmPFC and HPC during gradual attacks.

The current study also provides evidence that the activity in the amygdala is inversely coupled with the activity in vmPFC and HPC. The interaction between the frontal lobe and amygdala plays an important regulatory role under fear conditions. Previous research has highlighted the instructional role of the vmPFC-amygdala circuit in fear extinction [78, 79, 80], and these two regions are inversely coupled during the regulation of negative affect [81]. On the contrary, the amygdala may also inhibit the activation of the frontal lobe under fear conditions. A rodent study found prefrontal neurons reduce their spontaneous activity under aversive conditions and this depression is related to amygdala activity [82]. These findings provide evidence for the inversely coupled between the amygdala and PFC during fear-related processes. Our results build upon previous knowledge and provide empirical evidence demonstrating that this negative coupling also occurs in humans escaping under rapid attacks, with the direction from the amygdala to vmPFC and HPC. We interpret this phenomenon as the amygdala suppressing activation of the vmPFC and HPC under rapid attack. It may be attributed to the demanding nature of model-based planning, such as strategy making and memory retrieval, which consumes excessive computational resources and may not be advantageous for survival under immediate threat [4, 5]. Thus, the activity of the vmPFC and HPC may be inhibited by the reactive fear circuit, enabling prey to react quickly and ensure their survival and safety in the presence of rapid predators.

### Limitations and future work

Our study also has certain limitations and presents opportunities for future research. Due to limited electrode coverage and lack of a reliable high-resolution human brain atlas, our study did not subdivide the specific subregions within the targeted brain areas, such as the central nucleus or basolateral nucleus in the amygdala. Future research could employ a larger sEEG dataset and utilize high-resolution imaging techniques such as 7T fMRI [83] and a fine-grained human brain atlas [84] to investigate the role of more specific subregions in human escape behavior. Moreover, in comparison to the experimental paradigm, real-life escape scenarios faced by humans and animals are more complex. Future research should incorporate more naturalistic experimental paradigms, such as virtual reality simulations [85, 86, 87], to investigate escape decisions more realistically. Additionally, apart from the high gamma band, other frequency bands of brain signals, such as the theta band, may also play a role in escape [88]. Besides, other neural features such as EEG variability may also be associated with escape activity[89]. Future research could explore these additional features, and investigate the cross-frequency couplings between different brain regions, thereby providing a more comprehensive understanding of the neural dynamics underlying escape behavior. Furthermore, future research could also explore the use of brain stimulation techniques, such as transcranial alternating current stimulation (tACS) [90] or deep brain stimulation (DBS) [91], to establish causal relationships between the targeted brain circuit and human escape behavior. Such research in these directions can provide further insights into the neural basis of human escape behavior, and in characterizing adaptive and maladaptive decision-making under threat.

## Conclusion

In conclusion, our study utilized iEEG recordings and an FID task to examine the specific roles and interactions of two fear circuits across different stages of the escape process. We found that the cognitive fear circuit encoded the threatening types in the information process stage, while the reactive fear circuit mainly engaged in the escaping stage, particularly in response to rapid attacks. Furthermore, we observed that the amygdala actively suppresses the cognitive fear circuit during rapid attacks. These findings extend previous human functional magnetic resonance imaging results and animal evidence, revealing the distinct role of the reactive and cognitive fear circuits in coordinating the entire human escape process under different types of attacks.

## Methods

### Participants

We conducted intracranial recordings from a total of 18 patients suffering from drug-resistant focal epilepsy. Five patients were excluded from the analysis due to their limited cognitive ability or a lack of seriousness during the experiment, as evidenced by fewer than 15 correct trials in either condition. Additionally, one patient was excluded due to having undergone cortical resection surgery prior to the study. The demographic information of the remaining 12 subjects (5 females; age, 29.42 ± 10.04 years, mean ± SD) is shown in Table 1. All patients were implanted with stereoelectroencephalography (sEEG) depth electrodes for the localization of epileptogenic regions. The placement of the electrodes was solely based on clinical requirements. Each implanted electrode shaft had a diameter of 0.8 mm and contained 8-16 contacts along its length. Each contact measured 2 mm in length, with a distance of 3.5 mm between adjacent contacts. Both the Ethics Committee of the University of Macau (BSERE22-APP015-ICI-M1) and the Ethics Committee of the Hongai Hospital, where patient data were collected, approved the study. Additionally, written informed consent was obtained from all participants.

### Experimental paradigm

The current experimental paradigm was developed from the FID task [11]. Participants (blue circle) were instructed to imagine themselves as an animal (prey) and need to escape from the predator (triangle) (Figure 1 A, B, C). The participant and predator were positioned on a runway that stretched for 100 units in length, with the upper end of the runway designated as 0 units and the lower end as 100 units. In each trial, the prey foraging at a distance of 20 units above the safety zone located at the bottom of the runway. The predator would appear on the upper area of the runway and initially move slowly toward the prey at a speed of 20 units/s. Subsequently, the predator would accelerate to attack (speed up to 150 units/s) at a random distance, specific to the two predator types (fast vs. slow attacking). To successfully escape from the predator, participants needed to press the down arrow key on the keyboard before the predator accelerated. Failure to do so would result in the participant being captured. However, participants need to remain longer to get enough food under slow attack. Premature escape will result in a lack of physical strength to escape the predator’s future hunting. The boundary distance of the premature escape was determined by the Gaussian distribution with a mean value of 40 and a standard deviation of 3.

Our tasks have two types of predators: fast-attacking and slow-attacking (Figure 1 A). The difference between them lies in their different acceleration distances. The acceleration distance of the fast-attacking predator was determined by the Gaussian distribution with a mean value of 20 and a standard deviation of 3. The acceleration distance of the slow-attacking predator was determined by the Gaussian distribution with a mean value of 70 and a standard deviation of 3. There were 80 trials in the task, which were evenly divided into two blocks. We used two colors to represent two types of predators: purple and orange. We avoid using colors such as red or green that may contain safety or danger information. The predator–color relationship was randomly assigned and consistent within the same block. After the first block, the assignment of the–color relationship was altered to introduce novelty and avoid the (habitual) fixation of subject strategies.

In each trial, participants were first presented with a black cross on the screen for 2-3 s as visual fixation and baseline. Then, the cross will change its color to purple or orange, corresponding to the predator type as a cue of the predator type (0.5 s). Next, the predator starts to attack and participants need to make the escape decision for this trial and see the corresponding results. Under fast attacking conditions, the potential outcomes are captured and escaped. Under slow attacking conditions, the potential outcomes include captured, escaped, and premature escape (Figure 1 B). In the end, participants will receive reward visual stimuli for 2 s (happy images of animals) if they successfully escape. Or they will receive punishing stimuli (terrifying images of animals or images of animals being preyed on) if they are captured or prematurely escaped (Figure 1 C). After the trial, subjects are required to rate the difficulty of escape using a visual analog (1–5) scale.

### Intracranial EEG recordings, localization, and preprocessing

We used intracranial recordings in patients undergoing sEEG monitoring as part of clinical treatment for drug-resistant epilepsy. Electrophysiological signals were collected from electrodes implanted into the brain parenchyma with a 2 kHz sampling frequency. The signals were collected from standard clinical penetrating depth electrodes (Sinovation, China) implanted into the brain parenchyma. Each electrode is composed of 8–16 contacts, with a diameter of 0.8 mm for the depth electrode. The length of each contact was 2 mm, and the distance between contacts was 1.5 mm. Using the manufacturer’s ROSA planning software (version 2.5.8), the trajectory for all planned electrodes was prepared by loading and fusing MRI and CT scans. Then, preoperative planning MR images used with trajectory plans were merged with postoperative CT scans obtained with electrodes in place to assess the accuracy of the electrode position in relation to preoperative planned trajectories. For each patient, the number and placement of the electrodes were determined by a clinical team to localize epileptogenic brain regions as part of the patient’s evaluation for epilepsy surgery.

The electrode contacts were localized and anatomically labeled using the Brainstorm software [92] in MATLAB. The presurgical anatomical T1 MRI and the post-operative CT image with the sEEG electrodes were obtained for each patient. We first segmented the MRI using the CAT12 toolbox and warped the patient MRI to a standard MNI 152 template brain for group-level analyses using the SPM mutual information algorithm. Then, we co-registered the CT to the SPM12 toolbox (https://www.fil.ion.ucl.ac.uk/spm/software/spm12/) and located each sEEG electrode manually on the co-registered CT image (Figure S1). We used two parcellations to label each recording site. Using parcellation AAL3 [93], we labeled the electrodes in the insula, HPC, striatum (caudate, putamen, and pallidum), amygdala, and midcingulate cortex. Moreover, the electrodes in vmPFC were labeled by a vmPFC mask available at https://neurovault.org/images/18650/. We have also labeled the electrodes within PCC. However, the small number of available electrodes (only 4 electrodes in total from 2 subjects) prevented further analysis. In addition, PAG is located in the brainstem and cannot be implanted with electrodes.

sEEG data preprocessing was conducted using the EEGLAB toolbox in MATLAB. Raw sEEG data were manually checked by a neurologist or experienced data analyst for epileptic spiking and spread. Data in each channel with epileptiform or artifactual activity were excluded from further analyses. Preprocessing included resampling data from 2000 Hz to 1000 Hz, bandpass filtered using a Butterworth filter from 1-150 Hz, re-referenced (average reference across all non-noisy channels), and bandstop filtered at 50 and 100 Hz (Butterworth filter with 2 Hz bandwidth) to remove line noise and harmonics. In the end, data was epoched into trials that were time-locked to stimulus onsets, ranging from 0.5 s pre-stimulus to 4 s post-stimulus (−0.5 - 4 s epochs).

### High-gamma activity extraction

HGA is the broadband activity spatially precise measure of local neuronal population spiking [39, 33, 40]. We extracted the HGA using the MNE software [94] in Python. To extract high-gamma activity, we used a 10-cycle Morlet wavelet transform [95] to decompose the epoch data into 70-120 Hz [43, 44] spectra in 5 Hz bands. For within-ROI analysis, we averaged the time-frequency signals of all trials under the same conditions (e.g., fast escaped or fast captured) for each electrode. However, for functional connectivity analysis, we retained the time-frequency signals of each trial. Next, baseline correction is performed for each frequency band by dividing the power at all time points within a trial by the average baseline power before stimulation (−0.1-0 s) and taking the log (log-ratio) [96]. To reconstruct the one-dimensional HGA signal, within 70-120 Hz were averaged to produce a single HGA time course per electrode. Lastly, the HGA signal from each electrode was low-pass filtered with a Gaussian window (width = 0.1 s) for further analysis. After we extracted the HGA for Within-ROI analysis, for each condition, we obtained an HGA value matrix with dimensions of [Electrodes * Time points] across all participants. For the subsequent functional connectivity analysis, we obtained an HGA value matrix with dimensions of [Trials * Electrodes * Time points] for each condition in each participant.

### Within-ROI analysis

In our study, we conducted a within-ROI analysis to investigate the significance of neural activation between two conditions, as well as the differences compared with the baseline.To determine the significance between the HGA in the two conditions, we employed one-way ANOVA tests. To assess whether the HGA significantly deviated from the baseline, we utilized paired t-tests. To address the issue of multiple comparisons and ensure the validity of our results, we performed a cluster-based permutation test. This method takes into account the temporal proximity by clustering time series and is particularly suitable for analyzing time series data. We set the p-value threshold for permutation at 0.05 and conducted 10,000 permutations.

### Functional connectivity analysis

Functional connectivity (FC) between two ROIs was assessed by computing Pearson correlations between all inter-lobe electrode pairs. To estimate the directed FC, we employed the time-lagged cross-correlation analysis, as described in the study conducted by [45]. The lag range of cross-correlation is −300 to 300 ms, with a step size of 10 ms. Positive lags indicated that activity in ROI 1 preceded activity in ROI 2, while negative lags indicated the reverse, with activity in ROI 2 preceding ROI 1. A lag of zero indicated there is no delay between the two regions. As a result, a cross-correlation matrix was obtained for each condition with dimensions of [Trials * Electrodes pairs * Timelag].

For the averaged FC between two ROIs across all electrode pairs. We first averaged the trial dimension of the cross-correlation matrix and obtained a matrix for each condition with dimensions of [Electrodes pairs * Timelag]. We then employed the same statistical methods as in the Within-ROI analysis to investigate the significance of FC between the two conditions and its difference from the baseline. For each electrode pair, we extracted their FC matrix for each condition, with dimensions of [Trials * Timelag]. Using the same paired t-tests and permutation tests, we examined whether the FC of each electrode pair significantly differed from the baseline. Subsequently, for each significant cluster, we recorded the time lag and the FC value at the peak.

### Quantitative and statistical analysis

Paired t-tests were used to assess the statistical significance between the escaping behavior in the switching and non-switch conditions. Paired t-tests were also used to estimate the statistical significance between the HGA and the baseline. Pearson’s correlation was used to calculate the correlation between the HGA in different regions. One-way ANOVA was used to evaluate the statistical significance of the HGA or cross-correlation coefficient between different conditions. Cluster-based permutation tests [97] were utilized to calculate some statistics corrected for multiple comparisons using permutations and cluster-level correction.

## Acknowledgements

We are grateful to all the patients for participating in our study. The authors would like to thank Dr. Zhongzheng Fu for the early suggestions of the study. This work was mainly supported by the Science and Technology Development Fund (FDCT) of Macau [0127/2020/A3, 0041/2022/A], the Natural Science Foundation of Guangdong Province (2021A1515012509), Shenzhen-Hong Kong-Macao Science and Technology Innovation Project (Category C) (SGDX2020110309280100), MYRG of University of Macau (MYRG2022-00188-ICI) and the SRG of University of Macau (SRG2020-00027-ICI). We also thank all research assistants who provided general support in participant recruiting and data collection.

## Data and Code Availability

The sEEG data used in this study cannot be made publicly available but can be requested from the corresponding author [Haiyan Wu]. Code for the present data analyses is available at the GitHub repository: https://github.com/andlab-um/sEEG-slow-fast-attack.

## Supplementary Materials

**Figure S1:**
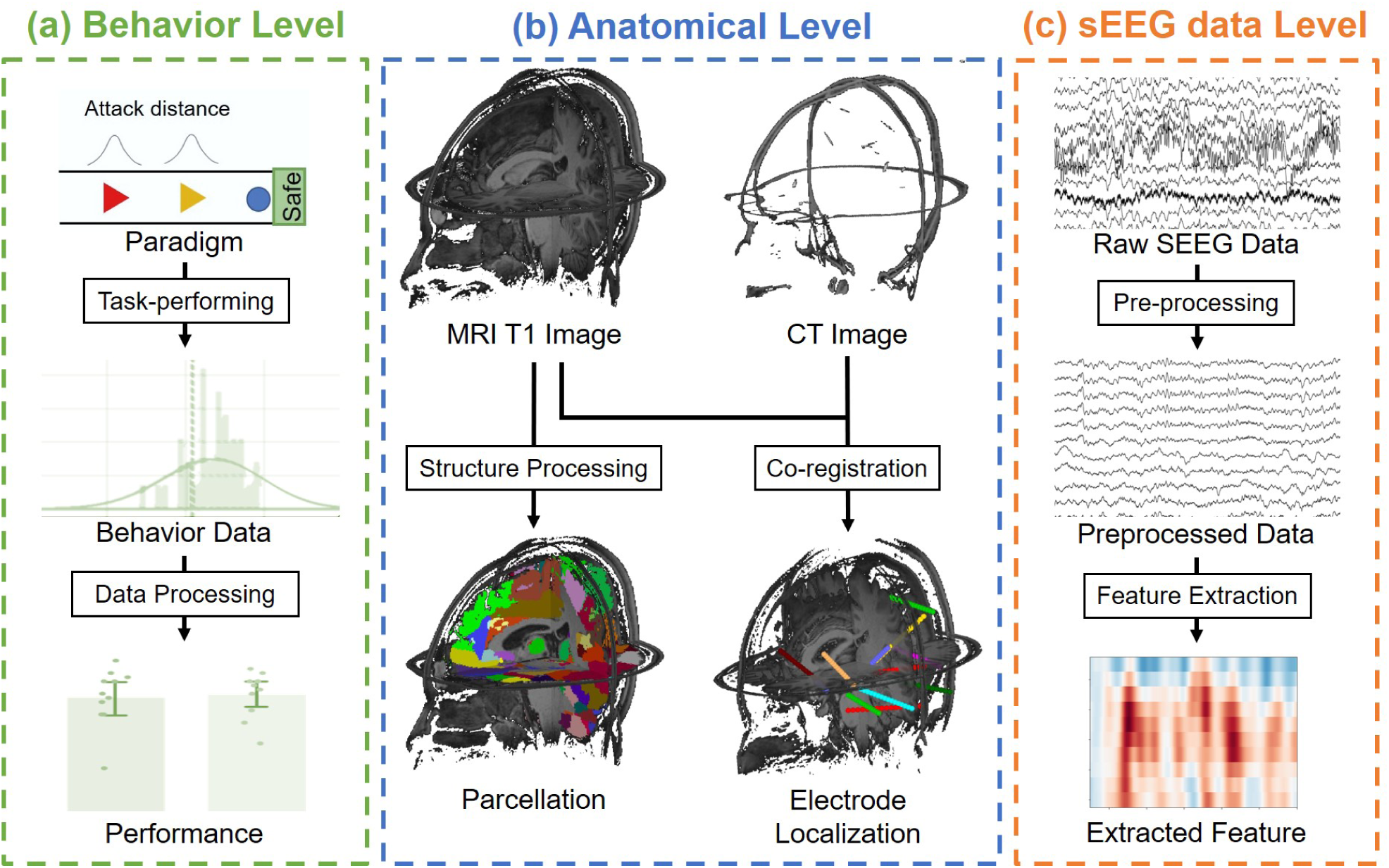
The data analysis process in the present study. (A) Behavior level. (B) Anatomical level. (C) sEEG level.

**Figure S2:**
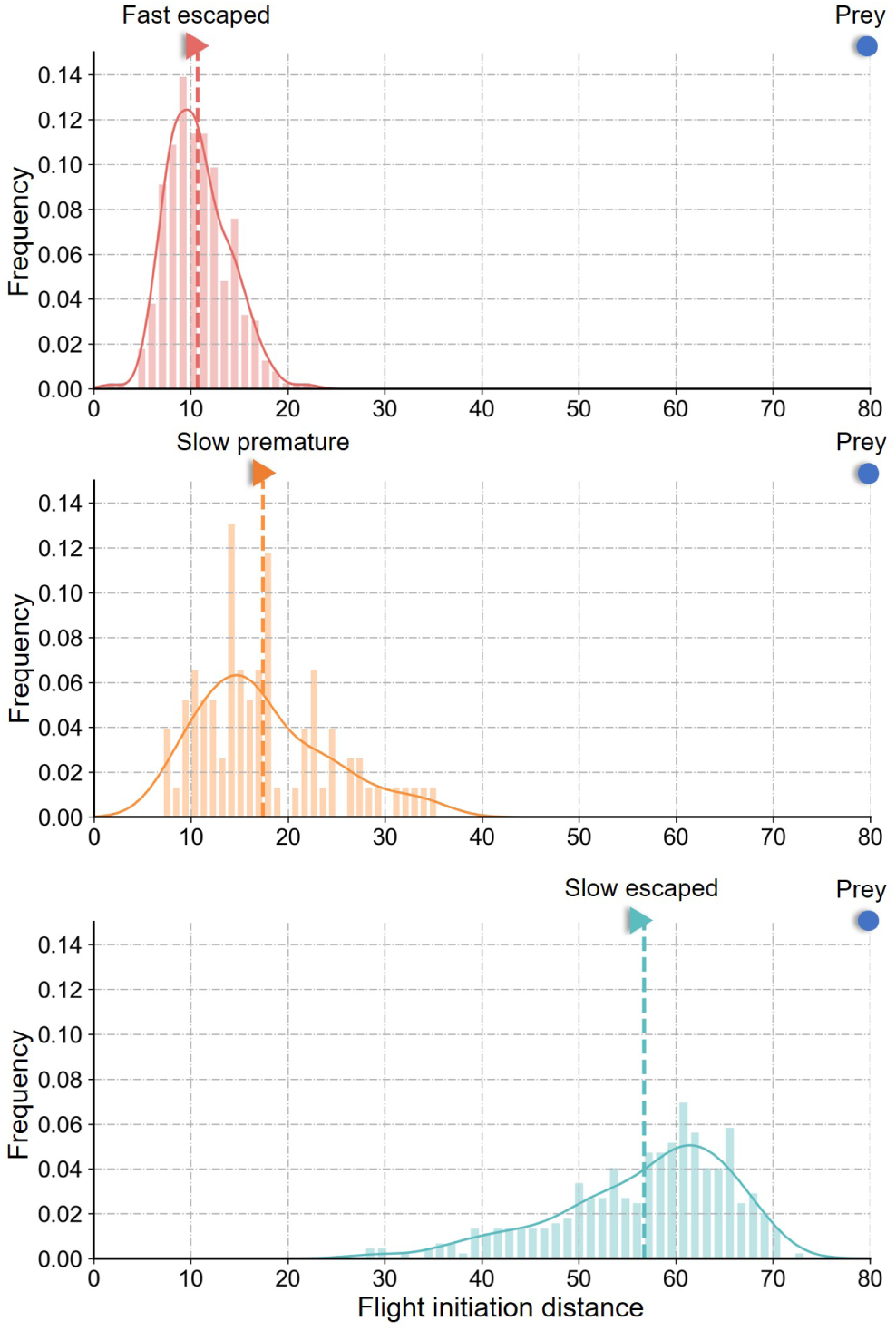
The distribution of the flight initiation distance (FID) in. (A) fast escaped trials, (B) slow premature trials, and (C) slow escaped trials.

**Figure S3:**
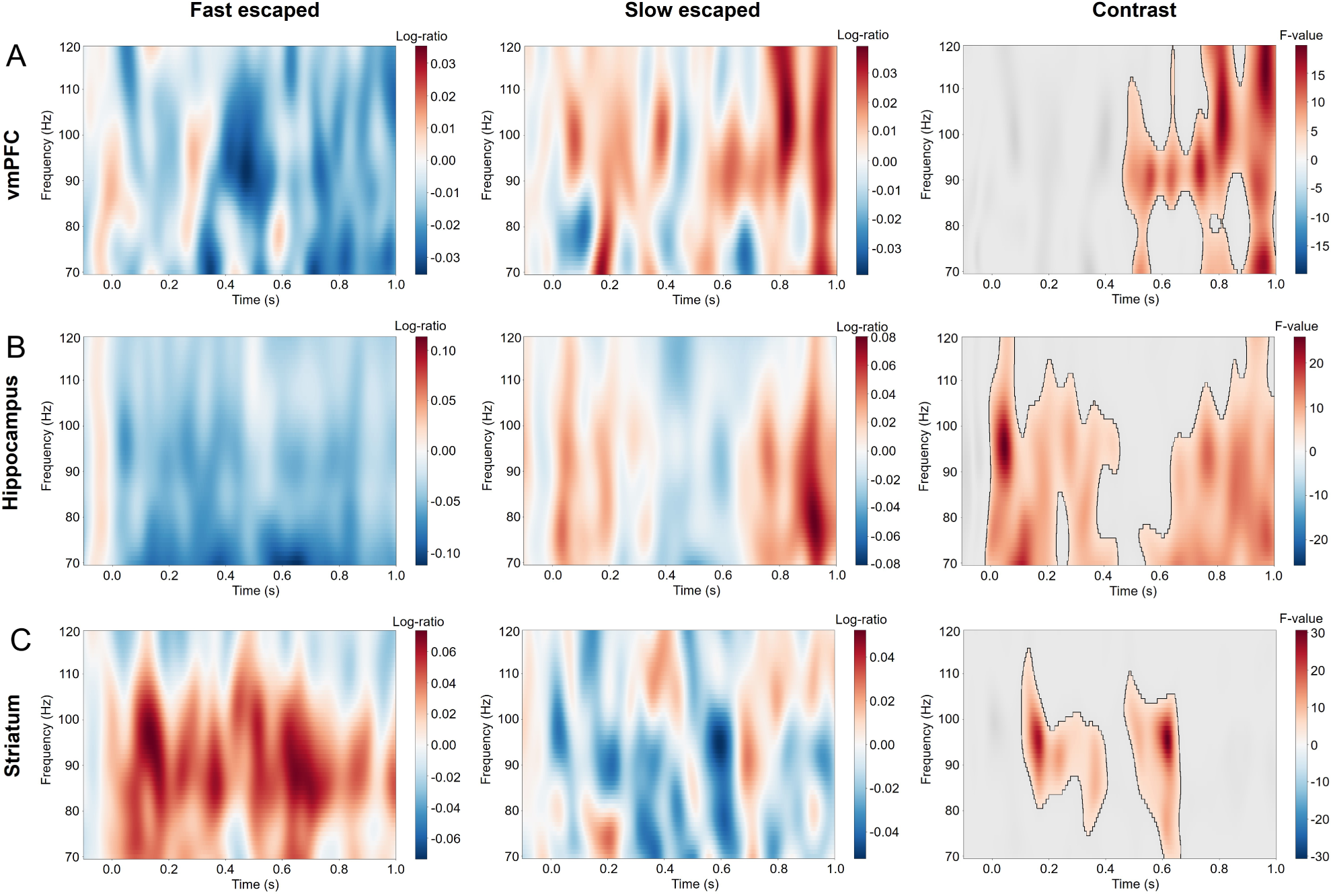
The time-frequency spectrograms under threat stimuli. The mean time-frequency spectrograms for all fast escaped trials (left), for slow escaped trials (middle), and contrast (Fast escaped versus slow escaped) with clusters after cluster-based permutation test for multiple comparison in (A) vmPFC, (B) hippocampus, and (C) striatum. The colorful region in the contrast map represents the significant time-frequency cluster that survive the cluster-based permutation test.

**Figure S4:**
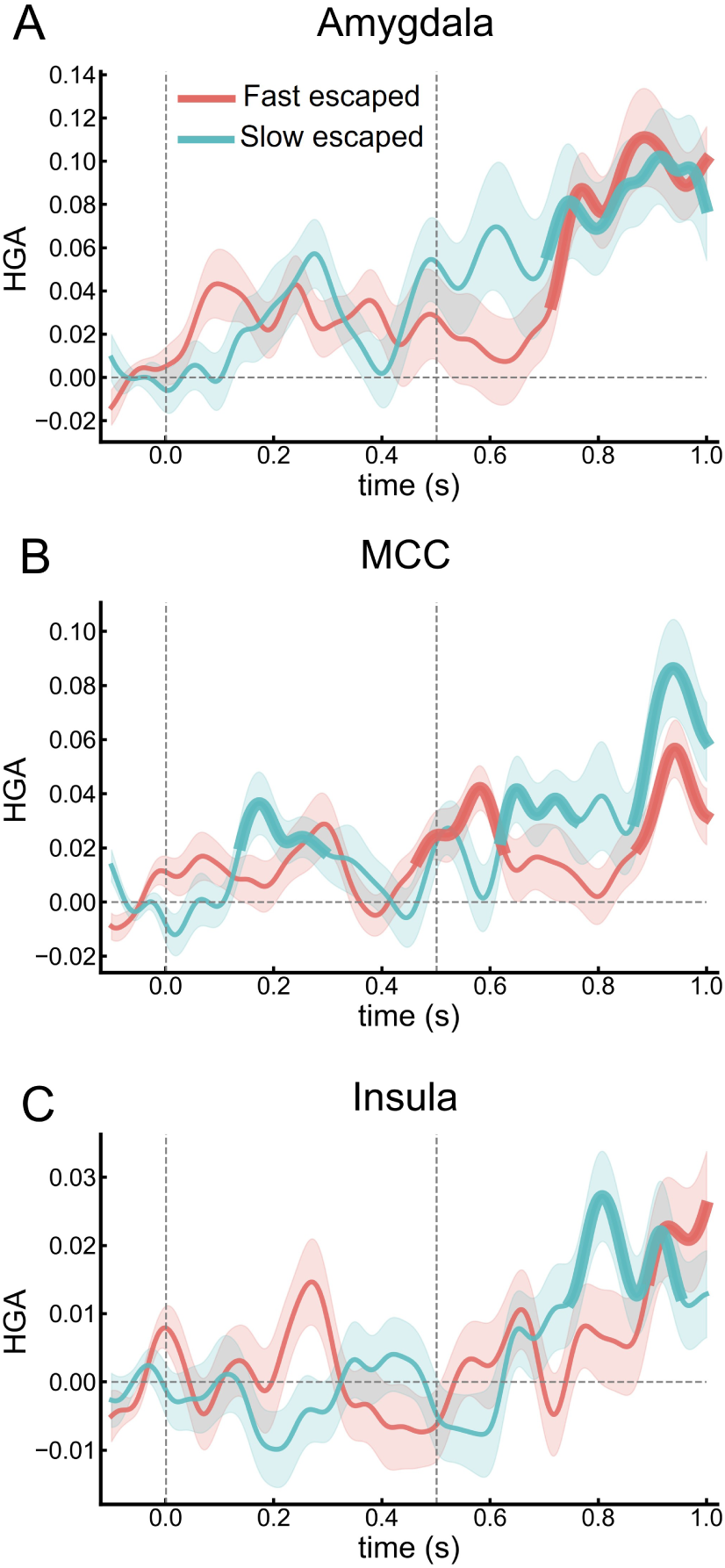
Intracerebral HGA under threat stimuli. The HGA under threat stimuli in 3 ROIs: (A) amygdala, (B) MCC, and (C) insula. The red and blue-green lines indicate HGA under fast escaped and slow escaped, respectively. The thicker line segment represents the period of significant difference between HGA and the baseline. The water-blue line below indicates the time period when there is a significant difference in HGA between the two conditions. Error bars represent the standard error of the mean HGA. The stimulus (cue of attacking condition) appears at 0 s in time series, and the attack begins at 0.5 s. In these three brain regions, there was no significant difference between the two conditions

**Figure S5:**
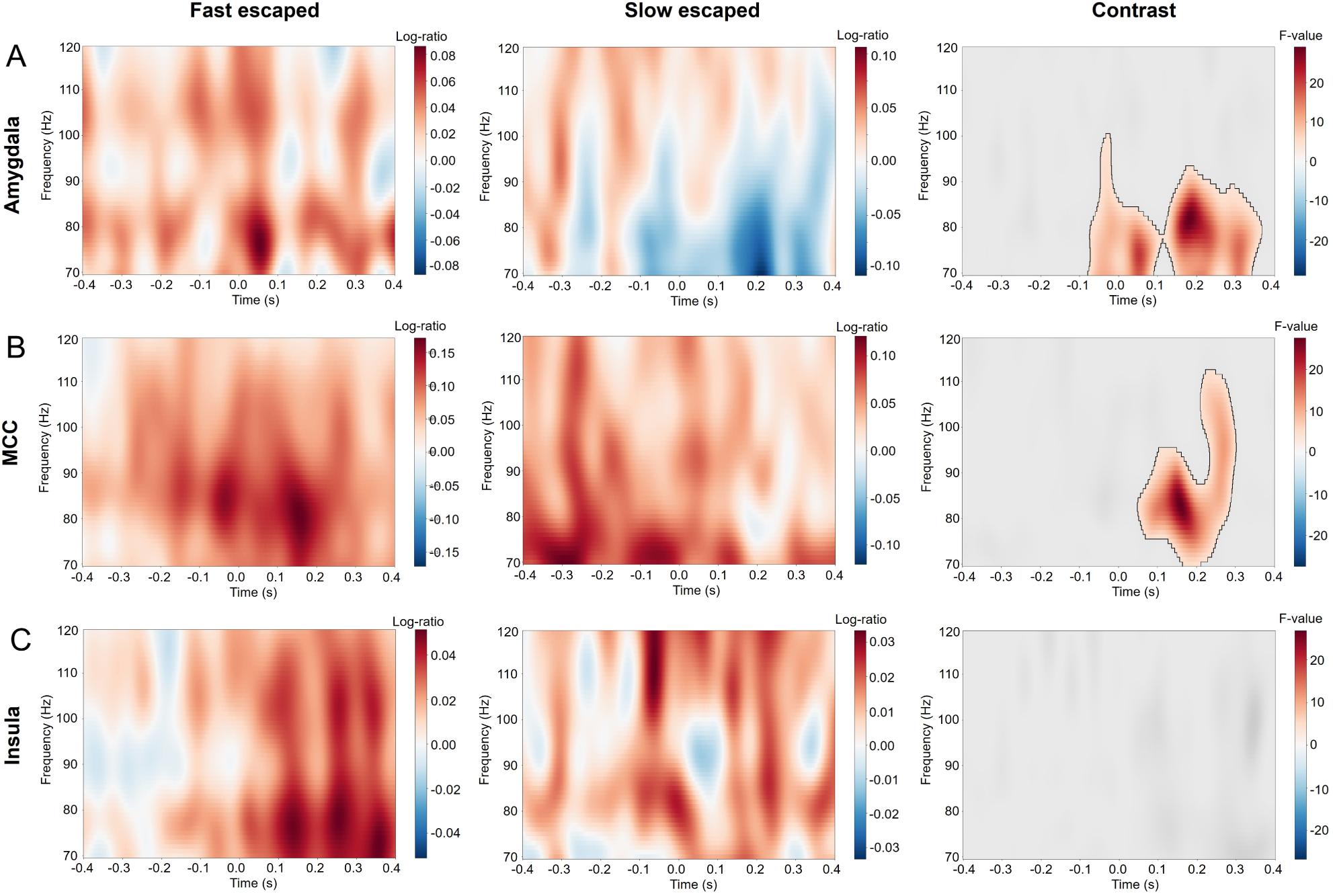
The time-frequency spectrograms in escaping. The mean time-frequency spectrograms for all fast escaped trials (left), for slow escaped trials (middle), and contrast (Fast escaped versus slow escaped) with clusters after cluster-based permutation test for multiple comparison in (A) amygdala, (B) MCC, and (C) insula. The colorful region in the contrast map represents the significant time-frequency cluster that survive the cluster-based permutation test.

**Figure S6:**
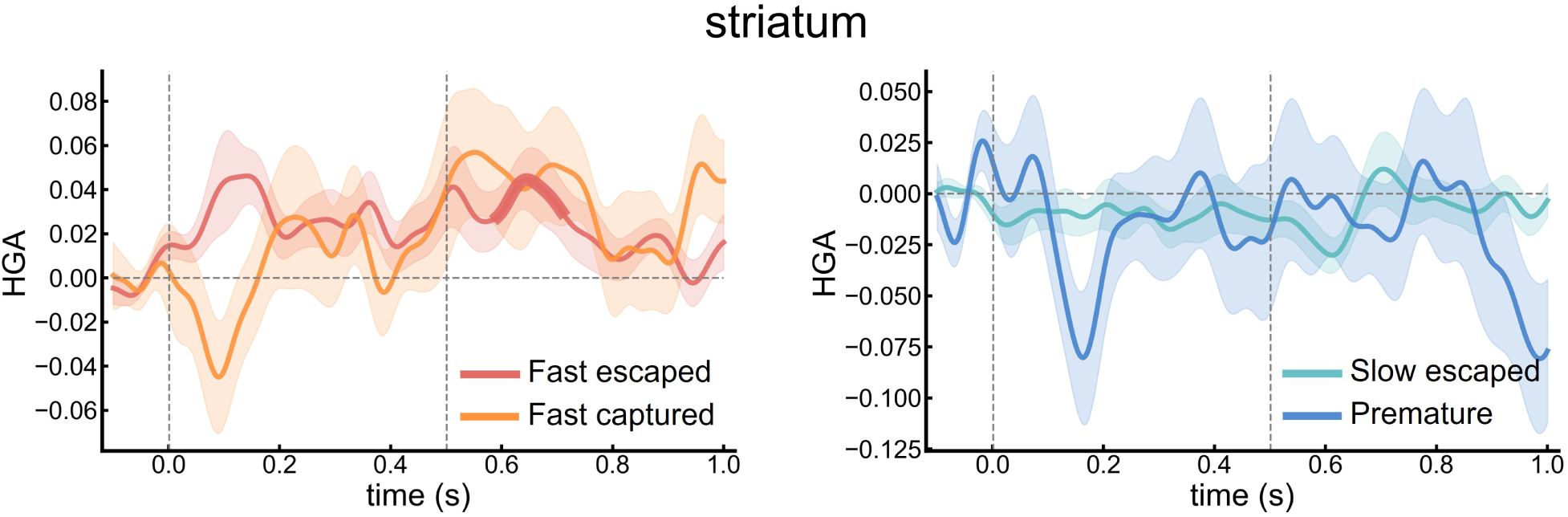
Intracerebral HGA between correct escape trials and failed escape trials. The HGA between (left) fast escaped trials and fast captured trials in striatum. The red and orange lines indicate HGA in the slow escaped trials and the slow premature escaped trials, respectively. The HGA between (right) slow correct escaped trials and the slow premature escaped trials in striatum. The blue-green and dark blue lines indicate HGA in the slow correct escaped trials and the slow premature escaped trials, respectively. In striatum, there was no significant difference between the successfully escaped trials or failed trails.

**Figure S7:**
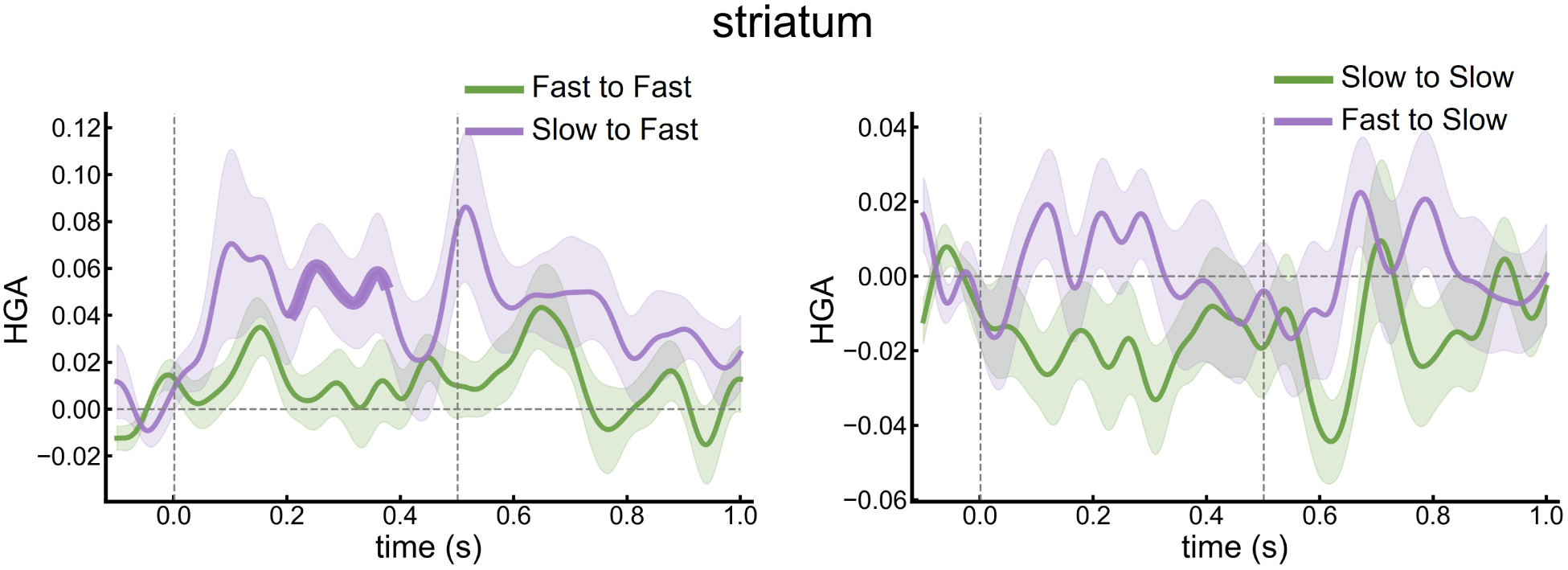
Neural activation of striatum under escaping condition switching. Left: The HGA between the fast-fast trials (green line) and slow-fast trials (purple line). Right: The HGA between slow-slow (green line) trials and fast-slow (purple line) trials. In striatum, there was no significant difference between the switching trials and no-switching trials.

**Table S1:** The number of electrode pairs between two brain regions in the functional connectivity analysis.

| SBJ | vmPFC-HPC | Amygdala-vmPFC | Amygdala-HPC |
| --- | --- | --- | --- |
| S01 | 60 | 6 | 10 |
| S02 | 65 | 15 | 39 |
| S05 | / | / | 48 |
| S06 | 60 | 60 | 16 |
| S07 | 140 | 100 | 35 |
| S08 | 336 | 168 | 98 |
| S09 | 195 | 234 | 30 |
| Total | 856 | 583 | 273 |

